# Sneezing in response to bright light exposure: A case study in a photic sneezer

**DOI:** 10.1101/2024.12.11.627890

**Authors:** Lucien Bickerstaff, Josef Trinkl, Stephan Munkwitz, Manuel Spitschan

## Abstract

**Background:** The photic sneeze reflex (PSR) is a widespread, yet understudied phenomenon characterised by sneezing in response to bright-light exposure, reportedly affecting around 30% of the general population. Our goals were to collect real-world data to characterise PSR-inducing naturalistic light conditions, and to develop an indoor protocol to reliably induce the PSR in affected individuals using parametric stimuli.

**Methods:** This study was carried out on one male adult affected by photic sneezing (n=1). To characterise naturalistic light conditions eliciting photic sneezing, real-world light exposure was measured over a 30-day period, while logging PSR events. To study photic sneezing in response to artificial stimuli, a setup including a multi-primary LED source and an integrating sphere was used to present 30-second light stimuli to the participant while collecting pupillometric data with an eye-tracker.

**Results:** 82 photic sneeze events were recorded, with an average of 2.73 sneezes per day and a range of 1 to 6 sneezes per event. At a sneeze event, illuminance is on average ten times bigger than five minutes before the sneeze event. A significant increase in illuminance is observed around 2 minutes before the sneeze event. Light levels go back down to pre-sneeze levels within 10 minutes after sneezing. Despite exposure to more than 150 stimuli, no sneeze could be artificially induced in the participant. However, a strong tickling sensation was consistently reported, especially for high illuminance settings.

**Conclusions:** Real-world light data confirmed that a sudden increase in environmental lighting conditions can induce photic sneezing. Further analysis could be relevant on instances of illuminance increments not eliciting a photic sneeze. The experimental setup only elicited tickling sensations, but with further testing and optimisation, it is reasonable to assume that it would reliably induce photic sneezes, thereby opening further mechanistic study of this intriguing phenomenon.

## Background

Photic sneezing is a widespread phenomenon, characterised by sneezing in response to bright light exposure (typically direct sunlight), reportedly affecting up to around 20-30% of the population (Askenasy, 1990; Dean, 2012; Everett, 1964; Kulas et al., 2017; McKusick, 2003; Morris, 1987). The photic sneeze reflex (PSR) has been documented for decades (Askenasy, 1990; Féré, 1890; Sédan, 1954), if not centuries, but despite its relatively high prevalence, is poorly understood. Some studies have attempted to further clarify the genetic and neural mechanisms involved in the reflex (Breitenbach et al., 1993; Chowdhury et al., 2019; Collie et al., 1978; Hydén & Arlinger, 2009; Langer et al., 2010; Peroutka & Peroutka, 1984; Sasayama et al., 2018), leading to no conclusive results.

To understand the naturalistic antecedents of the PSR, we examined sneezing in response to bright light exposure under naturalistic conditions while a photic sneezer logged their sneezes. In addition, we characterized tickle sensations and pupil responses to bright light exposure using white-light stimuli varying parametrically in illuminance.

## Methods

### Real-world light exposure measurements and photic sneeze logging

Real-life, daytime light measurements were carried out in summer between 12 July 2022 and 10 August 2022 (30 days) in and around Tübingen, Germany, as the participant went about his daily life. The participant, a healthy 21-year male (author LB of this manuscript), followed a Mon-Fri, 9-to-5 work schedule, working mostly indoors at a desk, at times with no natural light at all (only artificial light). Measurements were continuous from approximately 08:00 to 21:00, with a sampling frequency of 30 s. An ActTrust wearable actigraph with light logger (Condor Instruments, São Paolo, Brazil) was worn as a necklace so it would rest on the torso facing forwards. This would allow a good compromise between accurate measurements (i.e., in the same direction as the corneal plane) and practical comfort. The device was fixed to the torso using a magnetic plate resting between the participant’s torso and their clothing, to prevent unwanted tilt or flipping.

Every time the PSR manifested, the participant would self-report the sneeze event in a sneeze log datasheet implemented on Notion. Each entry contains the precise date and time of the sneeze (to the minute) and the number of sneezes for that event, since one sneeze event can contain multiple sneezes.

### Analysis of light exposure and PSR event data

The precise logging of each sneeze event allowed to obtain the average light exposure levels 20 min before and after the event. This was then compared to a reference – measured to be the average illuminance over 40-min, randomly-chosen time windows (n=100), when no sneeze event was reported. Individual contrasts between light exposure before and at each sneeze event were also analysed. Pre-sneeze light levels were averaged in time windows of 5 to 2 min before the sneeze event, and sneeze light levels in time windows ranging from 1 min before to 1 min after the sneeze event.

### Ethical approval

This case study was reviewed by the Ethics Committee of the Technical University of Munich (2024-74-W-SB). The participant gave informed consent.

### PSR induction in controlled conditions

A custom setup was used to elicit the PSR under controlled laboratory conditions. While wearing a head-mounted eye tracker for pupillometry (Pupil Core, Pupil Labs, Berlin, Germany), the participant was exposed to bright light emitted by a 10-primary light source (Spectra Tune Lab, Ledmotive, Barcelona, Spain) light into an integrating sphere.

Two paradigms were used to elicit the PSR:

1. A “one-shot” paradigm, where the participant would follow a 10-minute dark adaptation period, and then be exposed to a single light stimulus lasting 30 seconds;
2. And a 30-minute paradigm, where the protocol was identical but instead of one single stimulus, 24 stimuli were presented to the participant in succession, with 60 seconds of refractory darkness between the stimuli.

The photopic illuminance of the light stimulation varied between four different pre-set illuminance settings, measured at approximately 440, 1100, 4400 and 17600 lx (photopic) at eye-level using a calibrated spectroradiometer (STS-VIS, Ocean Optics, Ostfildern, Germany). An additional dark setting (0 lx) was included in the 30-min experiment. In addition to pupillometry, self-reports of sneeze onset (yes/no) and tickling sensation ratings from 0 (no tickle at all) to 10 (strong enough tickle to induce a sneeze) were obtained.

## Results

### Real-life light measurement and photic sneeze logging

Over the 30 day period, a total of 82 sneeze events were recorded, for an average of 2.73 sneezes per day. The number of sneezes range from 1 to 6 sneezes per event. Generally, a strong increase in light exposure can be observed in the few minutes leading to the sneeze event (Figure 1). Only one instance of photic sneezing was reported when light levels decreased before the sneezing event occurred. More specifically, a 10-fold increase in light intensity can be noticed just before photic sneezing on average, with quartiles within 2x to 30x. The difference in light intensity is significant in the 20 minutes surrounding the sneeze event. Pre-sneeze illuminance ranges from around 20 lx to 20 000 lx, while light levels at sneeze events range from around 1 000 lx to almost 30 000 lx (excluding the outlying instance mentioned just above, where illuminance fall to around 40 lx).

**Figure 1.**
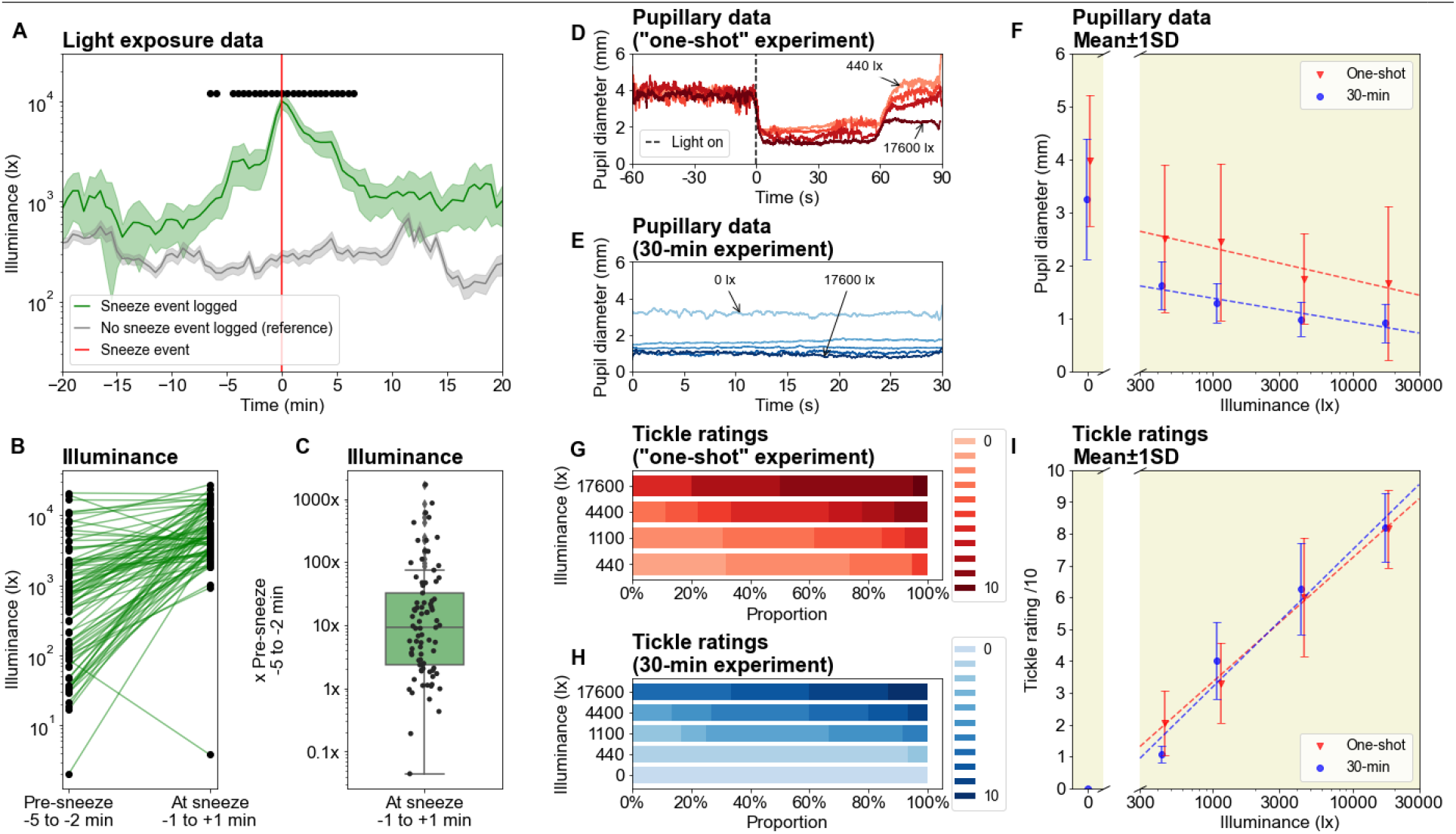
(A) Average light exposure 20 minutes before and after the sneeze event (n=82). Reference is the average light exposure over randomly-chosen 40-minute windows, in which no sneeze event was recorded (n=100). The light sensor was worn for 30 days during daytime (around 08:00 to 21:00), while all sneeze events were logged in parallel. Shaded error bars show standard error of the mean (SEM) and black dots show significance (independent t-test, p < .000610 after Bonferroni correction). (B, C) Difference between light exposure before (-5 to -2 min) and during (±1 min) sneeze events (linked t-test, t(81) = -8.21, p = 2.9e-12). (D) Average pupil diameter before and after light onset, for the one-shot paradigm. (E) Average pupil diameter during the 30-second light stimulations, for the 30-minute paradigm. (F) Comparison of the average pupil size as a function of illuminance for the two paradigms, mean ± STD. (G) Tickle ratings for the one-shot paradigm. (H) Tickle ratings for the 30-minute paradigm. (I) Comparison of the average tickle ratings as a function of illuminance for the two paradigms, mean ± STD.

After the sneeze event, the average illuminance typically falls back down to baseline levels within 10 minutes. The average light levels around photic sneezing events always remain above the reference, corresponding to the average illuminance for 100 time windows in which no photic sneeze was reported. Indeed, when sneezing was reported, the light levels almost always stayed above 500 lux, whereas when no sneezing was reported, they mostly remained below 500 lux. Sneezing often took place during transitions in environment, i.e., walking from home to the bus station, or from the bus station to the workplace.

### PSR induction in controlled conditions

In the experiments, we were unable to elicit a PSR using our laboratory stimuli. Despite exposure to more than 150 stimuli, the experimental setup could induce not sneeze in the participant. However, tickling sensations were consistently reported, and very high – although rare – values of 10/10 show that sneezing was very close under high light intensity. Both pupil and tickle ratings followed a monotonic relationship with the photopic illuminance of the stimuli.

## Conclusion

This case report provides a detailed characterization of photic sneezing in a known photic sneezer. Real-world data showed that sudden increases in illuminance often precede photic sneezing. These findings reinforce the connection between rapid light intensity changes and the PSR, while also suggesting thresholds of illuminance that may be critical for its manifestation. Despite the inability to artificially elicit sneezes in a controlled setting, the strong tickling sensations observed suggest that the experimental paradigm is promising with further refinement.

Future studies should investigate the variability in individual responses to artificial stimuli, as well as the underlying genetic, neurological, and sensory mechanisms of the PSR. Additionally, expanding the study to a larger cohort will be essential to understand phenotypic variation and interindividual differences in the PSR.

## References

Askenasy, J. J. (1990). The photic sneeze. Postgraduate Medical Journal, 66(781), 892–893. 10.1136/pgmj.66.781.892

Breitenbach, R. A., Swisher, P. K., Kim, M. K., & Patel, B. S. (1993). The photic sneeze reflex as a risk factor to combat pilots. Military Medicine, 158(12), 806–809.

Chowdhury, T., Sternberg, Z., Golanov, E., Gelpi, R., Rosemann, T., & Schaller, B. J. (2019). Photic Sneeze Reflex: Another Variant of the Trigeminocardiac Reflex? Future Neurology, 14(4), FNL32. 10.2217/fnl-2019-0007

Collie, W. R., Pagon, R. A., Hall, J. G., & Shokeir, M. H. (1978). ACHOO syndrome (autosomal dominant compelling helio-ophthalmic outburst syndrome). Birth Defects Original Article Series, 14(6B), 361–363.

Dean, L. (2012). ACHOO Syndrome. In V. M. Pratt, S. A. Scott, M. Pirmohamed, B. Esquivel, B. L. Kattman, & A. J. Malheiro (Eds.), Medical Genetics Summaries. National Center for Biotechnology Information (US). http://www.ncbi.nlm.nih.gov/books/NBK109193/

Everett, H. C. (1964). Sneezing in response to light. Neurology, 14(5), 483–483. 10.1212/WNL.14.5.483

Féré, C. (1890). Note sur l’éternuement provoqué par les excitations lumineuses. Comptes Rendus Des Séances de La Société de Biologie, 2(31), 555–557.

Hydén, D., & Arlinger, S. (2009). On light-induced sneezing. Eye, 23(11), 2112–2114. 10.1038/eye.2009.165

Kulas, P., Hecker, D., Schick, B., & Bozzato, A. (2017). Investigations on the prevalence of the photo-induced sneezing reflex in the German population, a representative cross-sectional study. European Archives of Oto-Rhino-Laryngology, 274(3), 1721–1725. 10.1007/s00405-016-4256-2

Langer, N., Beeli, G., & Jäncke, L. (2010). When the Sun Prickles Your Nose: An EEG Study Identifying Neural Bases of Photic Sneezing. PLoS ONE, 5(2), e9208. 10.1371/journal.pone.0009208

McKusick, V. A. (2003). ACHOO Syndrome. OMIM.

Morris, H. H. (1987). ACHOO syndrome. Prevalence and inheritance. Cleveland Clinic Journal of Medicine, 54(5), 431–433. 10.3949/ccjm.54.5.431

Peroutka, S. J., & Peroutka, L. A. (1984). Autosomal dominant transmission of the ‘photic sneeze reflex’. The New England Journal of Medicine, 310(9), 599–600.

Sasayama, D., Asano, S., Nogawa, S., Takahashi, S., Saito, K., & Kunugi, H. (2018). A genome-wide association study on photic sneeze syndrome in a Japanese population. Journal of Human Genetics, 63(6), 765–768. 10.1038/s10038-018-0441-z

Sédan, J. (1954). Réflexe photo-sternutatoire. Revue d’Oto-Neuro-Ophtalmologie, 26, 123–126.

